# Circadian regulation of human immunodeficiency virus type 1 replication

**DOI:** 10.1101/2022.11.30.518024

**Authors:** Helene Borrmann, Görkem Ulkar, Anna E. Kliszczak, Dini Ismed, Mirjam Schilling, Andrea Magri, James M. Harris, Peter Balfe, Sridhar Vasuvedan, Persephone Borrow, Xiaodong Zhuang, Jane A. McKeating

## Abstract

Human immunodeficiency virus 1 (HIV-1) is a life-threatening pathogen that still lacks curative therapies or vaccines. Circadian rhythms are endogenous daily oscillations that coordinate an organism’s response to its environment and invading pathogens. Recent reports of diurnal variation in peripheral viral load in HIV-1 infected subjects highlights the need for mechanistic studies. Here, we demonstrate rhythmic HIV-1 replication in circadian-synchronised model systems and show the circadian transcription factors, BMAL1 and REV-ERB, bind conserved motifs in the HIV-1 promoter. REV-ERB competes with ROR for binding to the same regulatory motif, and we uncover a role for ROR inhibitors to perturb rhythmic HIV-1 replication and host gene expression. We demonstrate that many HIV-1 host factors are circadian regulated and likely to define rhythmic viral replication. In summary, this study increases our understanding of the mechanisms underlying the circadian regulation of HIV that can inform HIV therapy and management.

## INTRODUCTION

HIV is the causative agent of acquired immunodeficiency syndrome (AIDS) and despite the increase in access to anti-viral treatments globally, there remains no cure. Integration of the viral genome into host chromatin is an essential step in the viral life cycle and infection is dependent on the host transcriptional machinery^1^. Viral transcription is regulated by the 5’ long terminal repeat (LTR) region that contains regulatory elements that recruit host transcriptional activators and repressors^2^. Current anti-retroviral therapies (ART) target multiple steps of the viral life cycle to suppress viral replication, but fail to eradicate long-lived reservoirs of viral integrants which perpetuate infection^3^. HIV-1 infects mainly haematopoietic CD4 expressing cells, that comprise the main site of virus replication, with T helper 17 (Th17) cells being particularly susceptible to infection^4^. Th17 cell differentiation depends on the master transcriptional regulator retinoic acid related-orphan receptor (RORC)^5^ that was recently reported to promote HIV-1 replication^6^, highlighting the importance of understanding the cellular transcriptional landscape in defining viral tropism.

The circadian clock is an endogenous timing system which oscillates in a 24 h manner coordinating many physiological processes. To anticipate environmental changes and minimise the risk of infection, immune functions depend on the cellular clock with many parameters oscillating throughout the day^7^. The outcome of infection by a wide range of pathogens is influenced by the time of day, and disruption of the circadian clock can increase disease severity^8^. On a molecular level, each cell has its own intrinsic clock machinery, which is orchestrated by several interlinked transcriptional/translational feedback loops^9^. The main circadian activators, brain and muscle ARNT-like 1 (BMAL1) and circadian locomotor output cycles kaput (CLOCK), form heterodimers that bind *cis* elements called E-boxes in the regulatory region of target genes which include their own repressors Period (*Per*) and Cryptochrome (*Cry*). An ancillary feedback loop is composed of the nuclear receptors REV-ERBs and RORs, themselves under transcriptional activation by CLOCK/BMAL1. REV-ERBs and RORs bind to ROR response elements (RORE) in the promoter regions of target genes, notably *Bmal1*, to inhibit or activate transcription, respectively^10^. Multiple small molecule modulators targeting core clock proteins (mainly CRYs, REV-ERBs or RORs) have been discovered and have potential for therapeutic avenues^11^. Additionally, optimizing the time of day for medicine and vaccine delivery can influence treatment efficacy and disease outcomes^12^.

Viruses are reliant on their hosts for replication, and recent reports demonstrate the role circadian systems play in viral infection^13-15^. Some of the first studies linking the circadian clock with HIV-1 were made two decades ago: Zeichner *et al* observed diurnal fluctuations of HIV-1 RNA in paediatric patient plasma^16^ and Ou *et al* reported conserved E-box motifs in the LTR^17^. We previously showed that REV-ERB agonists inhibit HIV-1 replication whilst antagonism or genetic silencing of REV-ERBs increased LTR activity^18^. Recent studies show an association between HIV-1 RNA and transcript levels of BMAL1^19^ and describe a rhythmic pattern of peripheral HIV-1 RNA in patients on ART^20^. These studies illustrate the clinical relevance of the interplay between HIV and the host circadian clock, however, there is limited evidence for circadian components directly regulating HIV replication and the mechanisms remain to be elucidated.

Here, we demonstrate that HIV-1 replication is rhythmic in an *in vitro* cellular system and provide evidence of BMAL1 and REV-ERBα binding the viral genome. Pharmacological inhibition of RORC using inverse agonists perturbed the cellular clock and inhibited HIV-1 transcription. We further show that host factors important for HIV-1 replication are clock regulated which can be perturbed by ROR modulation. Our work provides fundamental insights on the core circadian transcription factors and circadian-regulated host pathways that regulate HIV replication.

## RESULTS

### HIV-1 replication is rhythmic

To investigate a role for circadian pathways to regulate HIV-1 infection, we selected to use the human osteosarcoma cell line U-2 OS since they show robust oscillations *in vitro* following serum shock^21^ and are widely used in circadian research^22^. The single cycle HIV-1 reporter NL4.3 R-E-luc (NL4.3-luc) encodes luciferase as a readout for viral replication (**Fig.1a**) and provides a tool to quantify HIV-1 transcription without generating progeny virus and secondary reinfection events^23^. Complementing the glycoprotein-defective NL4.3-luc with vesicular stomatitis virus encoded G protein (VSV-G) generates pseudoparticles that can infect a broad range of cell types and bypasses the natural HIV entry receptors (CD4 and chemokine receptors)^24^. We show that U-2 OS cells support HIV-1 replication that requires integration into the host genome as evidenced by the antiviral activity of the integrase inhibitor raltegravir^25^ (**Supplementary Fig.1**). To assess whether HIV-1 replication is rhythmic, U-2 OS cells were infected with NL4.3-luc VSV-G and synchronised by serum shock. Cellular RNA was harvested at 4 h intervals to quantify HIV-1 Gag RNA by qPCR, and luciferase activity was measured at 30 min intervals (**Fig.1b**). We observed significant rhythmic HIV-1 replication over 48 h (**Fig.1c**) with a period of 24.7 h and peak viral replication at circadian time (CT) 12.2 h (p<0.00001, Fast Fourier Transform Non-linear Least Squares analysis FFT-NLLS^26^ in BioDare2^27^). Transfecting U-2 OS cells with a Bmal1 promoter-driven luminescence reporter (Bmal1-luc), enabled us to compare HIV replication and Bmal1 promoter activity. Of note, Bmal1 promoter activity and transcript levels displayed a similar period and peak expression to HIV-1 replication (**Fig.1d**). Synchronisation of U-2 OS cells was confirmed by assessing transcript levels of additional clock genes (**Supplementary Fig.2**). To extend these observations to a more physiological cell type, we attempted to synchronise a Jurkat CD4 T cell line and human primary CD4 T cells by serum shock, however, these cells failed to show robust circadian changes when measuring endogenous Bmal1 transcripts as previously reported, despite being rhythmic in circadian entrained mice and individuals^28^. In summary, we show that U-2 OS cells are a suitable *in vitro* model system to study HIV-1 infection and that viral transcription is rhythmic, confirming a role for the host circadian machinery in regulating HIV-1.

**Figure 1.**
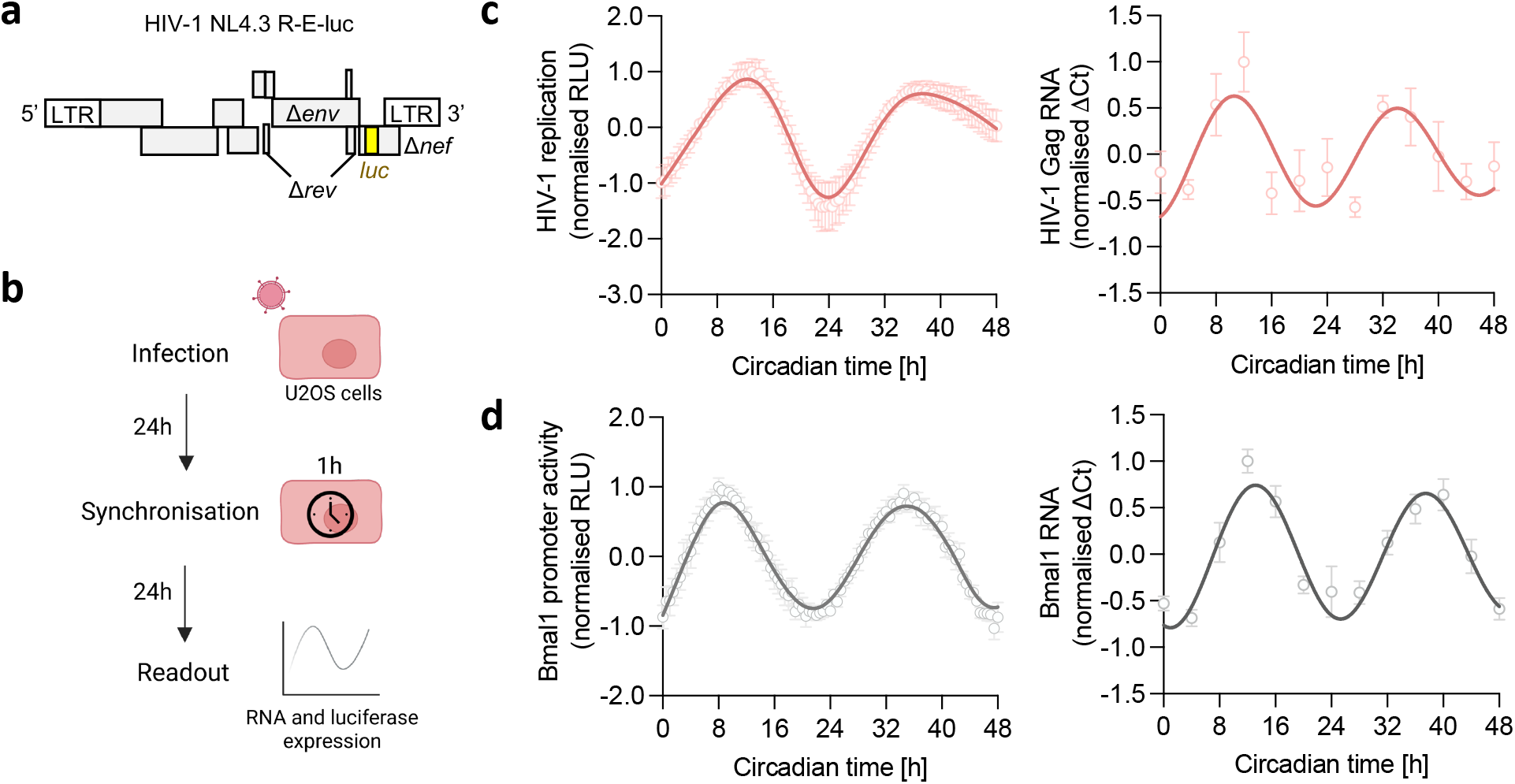
HIV-1 replication is rhythmic. **(a)** Cartoon of the HIV-1 NL4.3 R-E-luc (NL4.3-luc) reporter, encoding HIV genes with flanking long terminal repeats (LTR), including defective envelope (*Δenv*), regulator of expression of virion proteins (*Δrev*) and negative regulator factor (*Δnef*) and the luciferase (*luc*) gene which is the readout for viral replication. **(b)** U-2 OS cells were infected with HIV-1 NL4.3-luc VSV-G for 24 h followed by serum shock synchronisation for 1 h. 24 h later, luciferase activity was measured at 30 min intervals or cells harvested at 4 h intervals for RNA extraction for a total of 48 h. **(c)** U-2 OS cells were infected with HIV-1 NL4.3-luc VSV-G, synchronised and HIV-1 replication measured by luciferase activity (mean ± S.E.M., n=6) or cells were harvested at 4 h intervals for RNA extraction and HIV-1 Gag transcripts measured relative to a B2M housekeeper by qPCR (mean ± S.E.M., n=4). Analysis of luciferase data: eJTK cycle p<0.00001, period=24.7 h, peak expression=12.2 h (FFT-NLLS analysis, BioDare2). **(d)** U-2 OS cells stably expressing luciferase under control of the Bmal1 promoter (Bmal1-luc) were synchronised and promoter activity measured at 30 min intervals (mean ± S.E.M., n=7). Wild-type U-2 OS cells were synchronised, harvested at 4 h intervals, followed by RNA extraction and qPCR detection of Bmal1 RNA relative to a B2M housekeeper (mean ± S.E.M., n=4). All data are normalised to peak expression.

### BMAL1 regulates HIV-1 replication

Bmal1 is the only core clock gene whose deletion is sufficient to cause arrhythmicity of locomotor activity – the hallmark of circadian disruption^29^. Having noted a similar rhythmic pattern of Bmal1 and HIV-1 transcription, we assessed the role of BMAL1 in HIV infection. Rhythmic HIV-1 replication was blunted in stable Bmal1 knock-down (KD) cells compared to the parental control (**Fig.2a)**. Furthermore, silencing of Bmal1 or over-expression (OE) of BMAL1 and CLOCK in Jurkat cells resulted in a reduction or increase in HIV-1 replication compared to the control cells, respectively (**Fig.2b**). The HIV-1 LTR encodes four E-box motifs in the HIV NL4.3 strain (**Fig.2c**) and earlier reports showed that mutating these sites impaired HIV transcription^18,19^, supporting a model where BMAL1 binds and activates the HIV-LTR. To test this hypothesis, we isolated chromatin from NL4.3-luc VSV-G infected Jurkat cells and performed a BMAL1 chromatin immunoprecipitation (ChIP) assay. We designed primers to amplify the individual E-boxes in the LTR and showed enriched binding of BMAL1 to all motifs within the LTR, and to the *Per1* promoter compared to the IgG control (**Fig.2d**). Overall, these data provide evidence that BMAL1 binding to the HIV-LTR leads to enhanced HIV-1 transcription.

**Figure 2.**
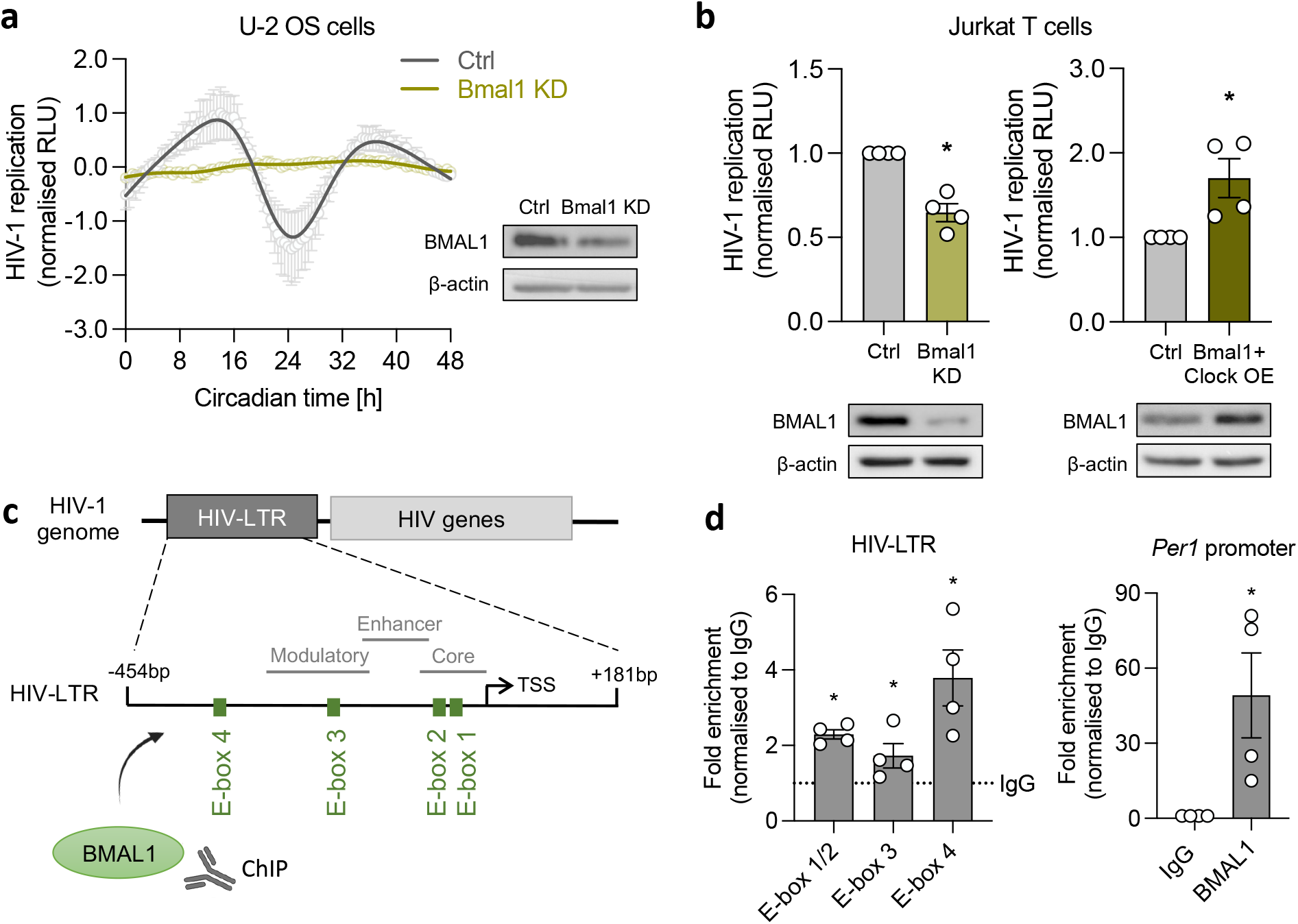
BMAL1 regulates HIV-1 replication. **(a)** U-2 OS parental control (ctrl) cells or U-2 OS Bmal1 knock-down (KD) cells generated by shRNA mediated silencing were infected with HIV-1 NL4.3-luc VSV-G. 24 h later, infected cells were synchronised by serum shock and viral replication measured by luciferase readout (mean ± S.E.M., n=3, normalised to peak, raw data in Supplementary Figure 7a). BMAL1 and β-actin protein levels were assessed by western blotting (representative of n=2). **(b)** Bmal1 KD or Bmal1/Clock over-expression (OE) in Jurkat cells was generated by siRNA mediated silencing (scrambled siRNA as ctrl) or transfection of plasmids (pcDNA3.1 as ctrl). Cells were infected with HIV-1 NL4.3-luc VSV-G and viral replication measured by luciferase activity (mean ± S.E.M., n=4, Mann-Whitney test, normalised to ctrl). BMAL1 and β-actin protein levels were assessed by western blotting (representative of n=2). **(c)** Cartoon of the 5’ HIV long terminal repeat (LTR) which encodes four E-box motifs (E-box 1: CAGCTG, E-box 2 & 3: CAGATG, Ebox 4: CAGTTG) in the NL4.3 strain, base pairs (bp) are shown relative to the transcriptional start site (TSS). **(d)** Binding of BMAL1 to the HIV-LTR was assessed by chromatin immunoprecipitation (ChIP). Jurkat cells were infected with NL4.3-luc VSV-G and chromatin extracts immunoprecipitated with BMAL1 antibody or rabbit IgG as a negative control. Fold enrichment of binding to either the E-boxes in the HIV-LTR or to the *Per1* promoter as a positive control was quantified by qPCR and is shown compared to non specific binding of IgG (mean ± S.E.M., n=4, Mann-Whitney test).

### ROR inhibition modulates host circadian clock factors and HIV-1 replication

Pharmacological agents targeting core circadian factors provide tools to study the mechanism(s) underlying the circadian regulation of HIV-1. While compounds directly targeting BMAL1 are not currently available, several drugs targeting RORs, regulators of *Bmal1* expression, have been developed^11^. Interestingly, the RORC isoform regulates Th17 cell differentiation^5^ and Salinas *et al* recently reported that the RORC inverse agonist GSK805 inhibits HIV-1 replication^6^. However, our knowledge of how ROR inverse agonists impact the cellular clock and rhythmicity of HIV replication is limited^11,30^. We confirmed that both U-2 OS and Jurkat cells express RORC (**Supplementary Fig.3a**). Treating synchronised U-2 OS cells with GSK805 showed a dose-dependent dampening of rhythmic Bmal1 promoter activity and endogenous transcript levels, with a 63.9% reduction in their amplitude at 10 µM compared to untreated cells (**Fig.3a**). We also noted a reduction in the rhythmic expression of several key circadian genes (*Rev-erbα, Per1, and Cry2*), all of which encode RORC binding motifs in their promoter regions (**Supplementary Fig.3b-c**). Reassuringly, GSK805 treatment showed a dose-dependent inhibition of Bmal1 promoter activity and protein expression in Jurkat cells (**Fig.3b**). To extend our observations to physiologically relevant cell types we treated CD8 depleted human peripheral blood mononuclear cells (PBMCs) with GSK805 and showed a reduction in *Bmal1, Rev-erbα, Per1*, and *Cry2* transcripts (**Fig.3c**), confirming a general perturbation of circadian gene expression.

**Figure 3.**
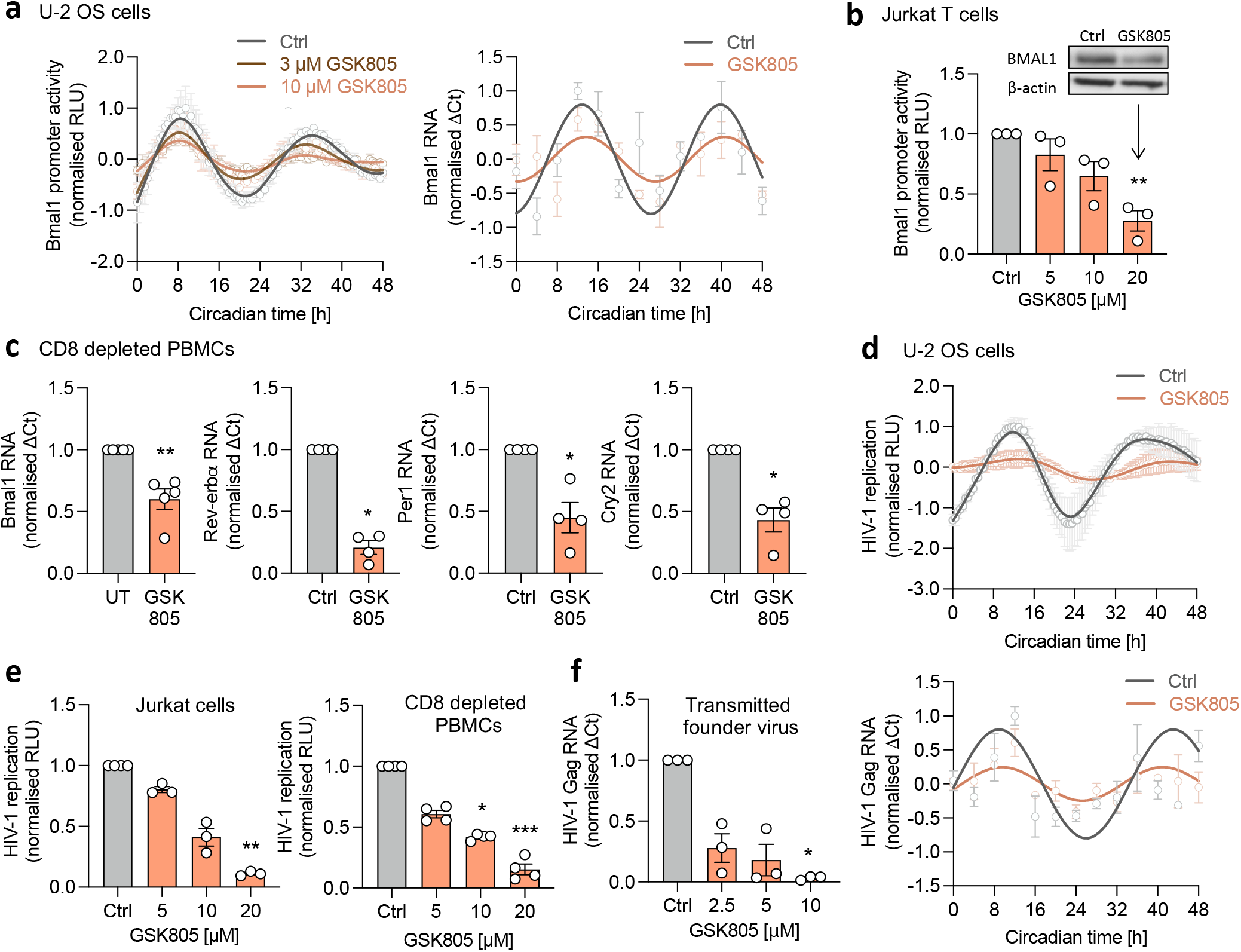
ROR inhibition modulates host circadian clock factors and HIV-1 replication. **(a)** U-2 OS cells stably expressing luciferase under control of the Bmal1 promoter (Bmal1-luc) were synchronised followed by treatment with GSK805. Luminescense was measured at 30 min intervals (mean ± S.E.M., n=3, raw data in Supplementary Figure 7b). Amplitude reduction compared to UT: 36.1% for 3 µM, 63.9% for 10 µM (FFT-NLLS analysis, BioDare2). Synchronised parental U-2 OS cells were treated with GSK805 (10 µM) and harvested at 4 h intervals, followed by RNA extraction and qPCR detection of Bmal1 transcripts relative to a B2M housekeeper (mean ± S.E.M., n=3). **(b)** Jurkat cells stably expressing Bmal1-luc were treated with GSK805 for 24 h and luciferase activity quantified (mean ± S.E.M., n=3, Kruskal-Wallis ANOVA). Jurkat cells were treated with GSK805 (20 µM) and Bmal1 and β-actin protein expression assessed by western blotting (representative of n=3). **(c)** CD8 depleted PBMCs were activated for 3 days with anti-CD3/CD28, treated with GSK805 (2.5 µM) for 7 days, followed by RNA extraction, and qPCR detection of Bmal1, Rev-erb⍺, Per1 or Cry2 RNA levels relative to a B2M housekeeper (mean ± S.E.M., n=4-5, Mann-Whitney test). **(d)** U-2 OS cells infected with NL4.3-luc VSV-G were synchronised, treated with GSK805 (10 µM) and luciferase activity measured at 30 min intervals (mean ± S.E.M., n=3, normalised to peak) or cells harvested at 4 h intervals, followed by qPCR detection of HIV-1 Gag RNA relative to a B2M housekeeper (mean ± S.E.M., n=3, raw data in Supplementary Figure 7c). **(e)** Jurkat cells or activated CD8 depleted PBMCs were infected with NL4.3-luc VSV-G, treated with GSK805 for 24 h and luciferase quantified as a readout of HIV-1 replication (mean ± S.E.M, n=3-4, Kruskal-Wallis ANOVA). **(f)** Activated CD8 depleted PBMCs were spinoculated for 2 h with patient derived HIV-1 (transmitted founder virus clone CH185) and cultured with GSK805 for 7 days, followed by qPCR detection of HIV-1 Gag RNA relative to B2M housekeeper (mean ± S.E.M., n=3, Kruskal-Wallis ANOVA). All data are expressed relative to UT control.

Intrigued by our observations of cellular circadian disruption by GSK805 we hypothesised inhibition of rhythmic viral replication. GSK805 treatment dampened the rhythmic expression of HIV replication and viral RNA in synchronised U-2 OS cells (**Fig.3d**). We confirmed GSK805 inhibition of HIV replication in Jurkat and CD8 depleted PBMCs (**Fig.3e**). To extend these observations we used a patient-derived founder strain of HIV-1 (CH185) (**Supplementary Fig.3d**) and GSK805 treatment significantly reduced viral transcript levels (**Fig.3f**). Of note, the compound showed no cytotoxicity at the concentrations tested in all cell types **(Supplementary Fig.4)**.

To ascertain whether a chemically different ROR inhibitor shows the same activity we studied the RORC inhibitor GSK2981278, that was evaluated in clinical trials as a topical agent for the treatment of psoriasis^31^. GSK2981278 treatment inhibited rhythmic Bmal1 promoter activity in U-2 OS, showing an 88.9% reduction in amplitude at 40 µM compared to untreated cells (**Fig.4a**). We also observed a dose-dependent reduction in Bmal1 promoter activity and protein expression in Jurkat cells (**Fig.4b**), and reduced Bmal1 transcript levels in CD8 depleted PBMCs (**Fig.4c**), with no evidence for cytotoxicity (**Supplementary Fig.5**). Importantly, GSK2981278 inhibited the replication of HIV strains NL4.3 and CH185 in Jurkat and primary cells (**Fig.4d-e**). Collectively, these data show that pharmacologically targeting RORC disrupts the cellular clock and inhibits rhythmic HIV-1 replication.

**Figure 4.**
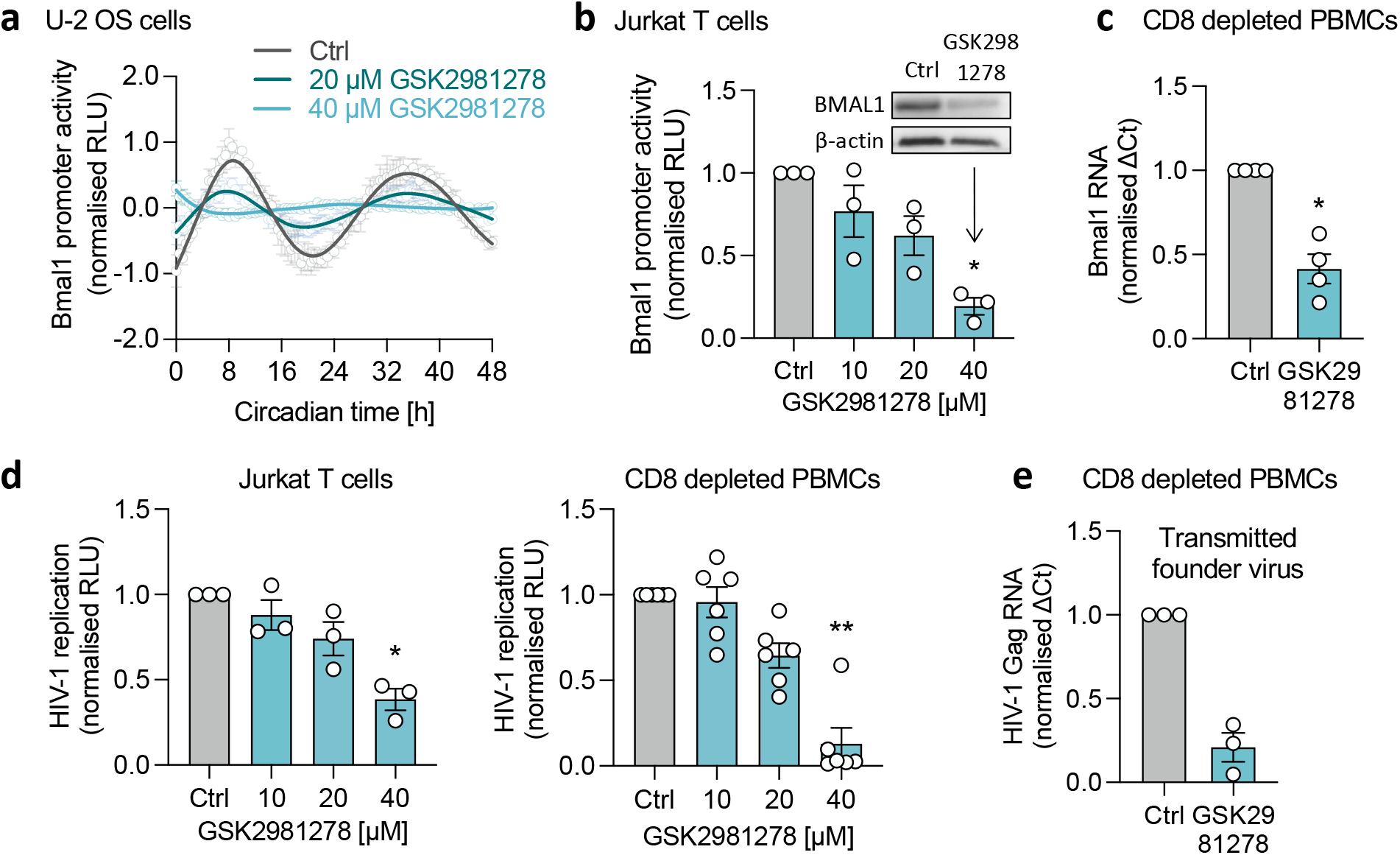
An independent ROR inhibitor reduces Bmal1 expression and HIV-1 replication. **(a)** U-2 OS cells stably expressing luciferase under control of the Bmal1 promoter (Bmal1-luc) were synchronised, treated with GSK2981278 and luciferase activity measured at 30 min intervals (mean ± S.E.M., n=4, raw data in Supplementary Figure 7d). Amplitude reduction compared to UT: 63.9% for 20 µM, 88.9% for 40 µM (FFT-NLLS analysis, BioDare2). **(b)** Jurkat cells stably expressing Bmal1-luc were treated with indicated concentrations of GSK2981278 for 24 h and luciferase activity quantified (mean ± S.E.M., n=3, Kruskal-Wallis ANOVA). Wild-type Jurkat cells were treated with GSK2981278 (40 µM) and BMAL1 and β-actin protein expression assessed by western blotting (representative of n=3). **(c)** CD8 depleted PBMCs were activated for 3 days with anti-CD3/CD28, cultured in the presence of GSK2981278 (20 µM) for 7 days, followed by RNA extraction and qPCR detection of Bmal1 RNA relative to a B2M housekeeper (mean ± S.E.M., n=4, Mann-Whitney test). **(d)** Jurkat cells (mean ± S.E.M., n=3, Kruskal-Wallis ANOVA) or activated CD8 depleted PBMCs (mean ± S.E.M., n=6, Kruskal-Wallis ANOVA) were infected with NL4.3-luc VSV-G, treated with different doses of GSK805 for 24 h and luciferase quantified as readout of HIV-1 replication. **(e)** Activated CD8 depleted PBMCs were spinoculated for 2 h with patient derived HIV-1 (transmitted founder virus clone CH185), cultured in medium with GSK2981278 (20 µM) for 7 days, followed by RNA extraction, and qPCR detection of HIV-1 Gag RNA relative to a B2M housekeeper (mean ± S.E.M., n=3). All data are expressed relative to UT control.

### REV-ERBα binds a conserved RORE in the HIV-LTR

RORC has been shown to regulate HIV replication directly through binding to a RORE in the HIV-LTR ^6^. Analysis of the RORE motif in LTR sequences deposited in the Los Alamos Database shows a low level of variation (**Fig.5a**), therefore we screened a panel of HIV LTR-Luc reporter plasmids encoding the major HIV-1 clades A-G [56] for their response to ROR inhibition. All of the HIV-LTR reporters were sensitive to ROR inverse agonist treatment (**Fig.5b**), consistent with the conservation of the RORE motif in these representative viral strains^18^. Since ROR often competes with the circadian repressor REV-ERB for binding to ROREs^32^, we hypothesised that REV-ERB would bind the HIV-LTR. To test this, we isolated chromatin from HIV-1 infected Jurkat cells (**Fig.5c**), performed a ChIP assay and observed REV-ERBα binding to the RORE in the HIV-LTR and its host target Bmal1 promoter (**Fig.5d**). Importantly, ROR inhibition by the inverse agonist GSK805 increased the binding of REV-ERBα to the HIV-LTR (**Fig.5e**), supporting our conclusion that RORC and REV-ERBα compete to regulate HIV transcription through the RORE element.

**Figure 5.**
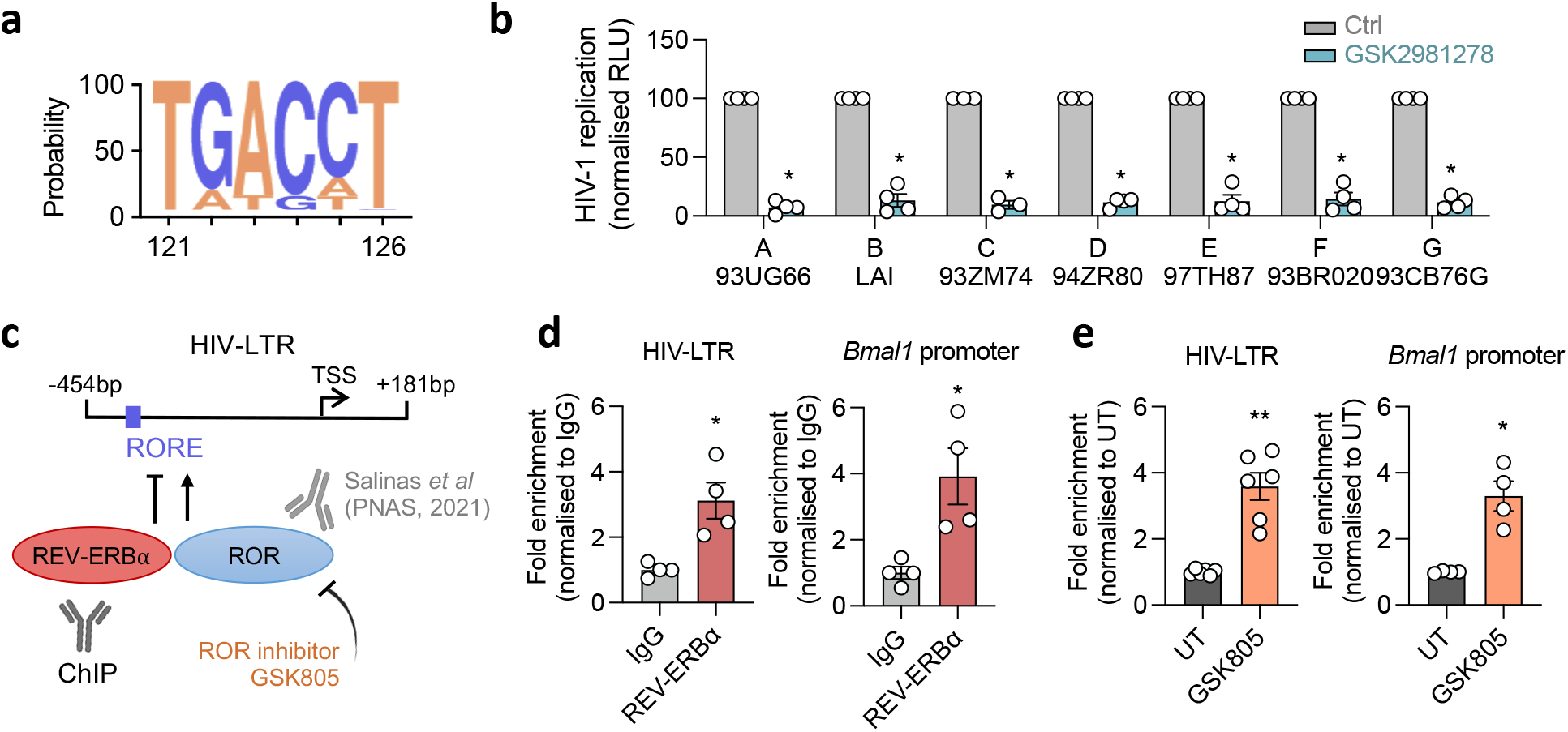
REV-ERB⍺ binds a conserved RORE in the HIV-LTR. **(a)** Consensus plot showing the conservation of nucleotides in position 121 to 126 of 1266 HIV-1 sequences deposited in the Los Alamos Database (coordinates are from the HXB2 reference). Bases of coding strand are displayed which correspond to the nucleotides ‘AGGTCA’ on the antisense strand, thereby representing the ROR response element (RORE) ‘RGGTCA’. Conservation is reflected by the height of the bases (y axis 0–100%). **(b)** Jurkat cells were transfected with reporter constructs of HIV-1 subtypes A-G, followed by GSK981278 treatment (40 μM) and LTR activity measured by quantifying luciferase activity 24h later (mean ± S.E.M., n=4, Mann-Whitney test). **(c)** Location of the RORE in the 5’ HIV-LTR of NL4.3 strain, base pairs (bp) are shown relative to the transcriptional start site (TSS). Model where ROR and REV-ERB⍺ compete for binding to the HIV-LTR which can be tested by chromatin immunoprecipitation (ChIP). RORC binding to the HIV-LTR was shown by Salinas *et al*^6^. **(d)** Jurkat cells were infected with NL4.3-luc VSV-G for 24 h and chromatin extracts immunoprecipitated with anti-REV-ERB⍺ or rabbit IgG as a negative control. Fold enrichment of binding to the RORE in either the HIV-LTR or the *Bmal1* promoter was quantified by qPCR and is shown compared to the IgG control (mean ± S.E.M., n=4, Mann-Whitney test). **(e)** Jurkat cells were infected with NL4.3-luc VSV-G followed by treatment with GSK805 (10 µM) for 24 h. Fold enrichment of REV-ERB⍺ binding to the HIV-LTR RORE or the *Bmal1* promoter was compared between untreated and GSK805 treated cells (mean ± S.E.M., n=4-6, Mann-Whitney test). All data are expressed relative to the control cells.

### HIV-1 host factors are regulated by the circadian clock

HIV-1 replication is dependent on a myriad of host factors, many of which could be affected by circadian rhythms. Several studies have identified cellular proteins that positively (dependency factors) or negatively (restriction factors) influence HIV replication^33,34^. Using a high-throughput CRISPR–Cas9 gene editing approach, Hiatt *et al* recently identified 90 genes that regulate HIV replication in human CD4 T cells^35^. To estimate whether transcript levels of these host factors are rhythmic we interrogated the Circa Database^36^ and showed that 43% of these genes are cycling across human tissues (**Fig.6a, Supplementary Fig.6a**). To extend this analysis, we examined several experimentally validated ChIP-seq data sets: including BMAL1 regulated genes from the mouse liver^37^ and REV-ERB^38^ or RORC^39^ regulated genes from murine Th17 cells. Comparing the BMAL1, REV-ERBα and RORC regulated target genes with the 90 HIV host factors^35^ showed 28 BMAL1, 31 REV-ERBα and 60 RORC regulated HIV host factors **(Fig.6b, Supplementary Fig.6b-c**). To identify HIV-1 associated host factors which are likely to be under direct BMAL1, REV-ERB or ROR control, we analysed their promoter regions for the presence of circadian motifs using HOMER (Hypergeometric Optimization of Motif EnRichment tool^40^). Our analysis identified 35 genes encoding a RORE, and 74 genes with an E-box (CANNTG), with 10 having the canonical E-box motif (CACGTG, **Fig.6c**). Notably, 5 promoters encoded both RORE and E-box motifs **(Supplementary Fig.6d)** giving a total of 40 genes. To validate this bioinformatic analysis, we screened all 40 genes for their response to RORC inverse agonists treatment in Jurkat cells. Seven genes were identified where GSK805 and GSK2981278 significantly reduced their expression (**Fig.6d**), consistent with their antiviral activity. Gene ontology (GO) cellular components analysis revealed that these 7 genes are involved in multiple cellular processes, ranging from transcriptional control to protein transport (**Fig.6e**)^41^. In summary, we have identified circadian regulated host factors which potentially contribute to rhythmic HIV-1 replication and can be modulated by ROR inhibition.

**Figure 6.**
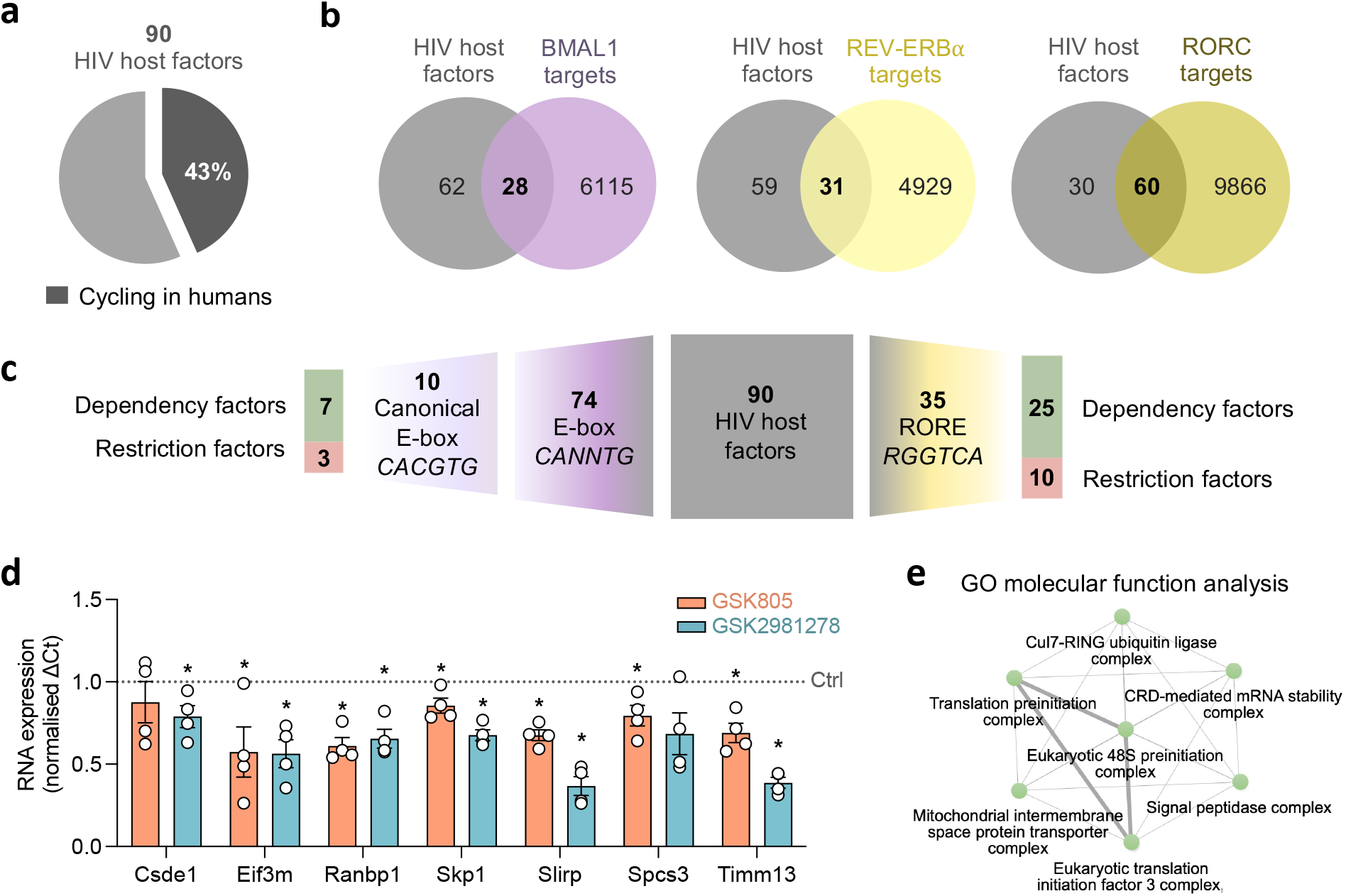
HIV-1 host factors are regulated by the circadian clock. **(a)** Expression of 90 HIV-1 host proteins^35^ was analysed using the CircaDatabase^36^ and 43% of genes identified as cycling in humans. **(b)** BMAL1 regulated genes^37^, REV-ERB regulated genes^38^ and RORC regulated genes^39^ were compared with host factors known to alter HIV-1 replication^35^. **(c)** HOMER (Hypergeometric Optimization of Motif EnRichment tool^40^) was used to analyse −1kb promoter regions of HIV-1 host factors and identified gene promoters encoding E-box motifs or ROR response elements (RORE). **(d)** Jurkat cells were treated with GSK2981278 (40 μM) or GSK805 (20 μM) for 24 h, cells were lysed, RNA was extracted, and gene expression analysed via qPCR (mean ± S.E.M., n=4, Mann-Whitney test). **(e)** Gene ontology (GO) cellular components analysis^41^ where each node represents an enriched GO term. Related GO terms are connected by lines, where thickness reflects the percentage of overlapping genes. See related Supplementary Figure 6.

## DISCUSSION

In this study we show that HIV-1 replication is rhythmic in circadian synchronised cells and identify a direct role for BMAL1 to activate viral replication via binding to the LTR. Pharmacological perturbation of circadian cycling by ROR inhibition dampened rhythmic HIV-1 replication and inhibited the activity of LTR sequences from diverse clades, suggesting that targeting host circadian components may be a pan-genotypic antiviral strategy. Further we provide evidence that REV-ERBα binds to a conserved RORE in the viral regulatory region and this binding was enhanced by ROR inhibition, suggesting competition between these nuclear receptors for binding to the HIV-LTR, similar to their competition for *Bmal1* and other clock-controlled genes^32^. Finally, we identify and validate circadian regulated HIV-1 host factors which may contribute to rhythmic HIV-1 replication **(Fig.7)**.

**Figure 7.**
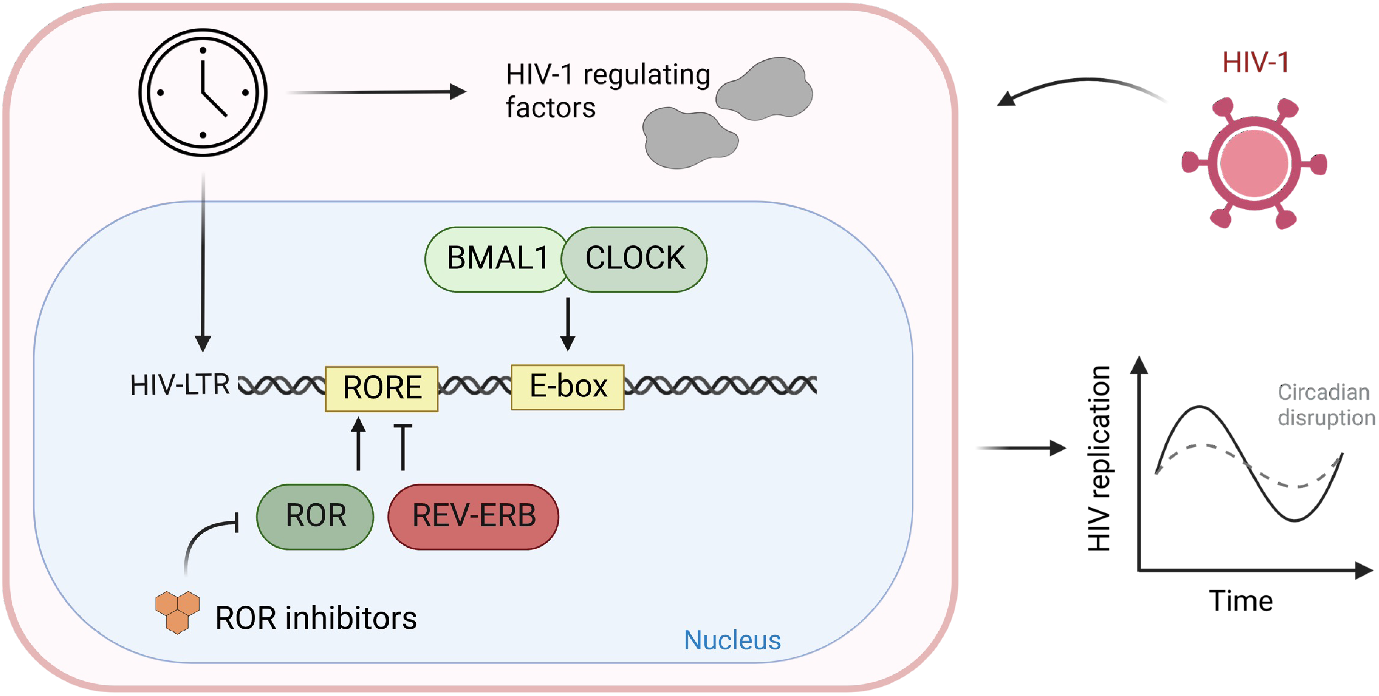
Model for circadian regulation of HIV-1. HIV-1 replication is dependent on circadian regulated host pathways. BMAL1 binds to E-boxes while REV-ERB⍺ competes with RORC for binding to the cognate ROR response element (RORE) in the HIV-1 long terminal repeat (LTR). RORC inhibitors disturb the cellular circadian rhythm that limit HIV-1 replication and increase REV-ERB⍺ binding to the HIV-LTR. The cellular clock regulates host factors for HIV replication that may indirectly contribute to the circadian regulation of HIV-1.

We observed that BMAL1/CLOCK over-expression increased HIV-1 replication in T cells, supporting a positive role for BMAL1 in driving HIV-1 transcription, consistent with an earlier report showing increased HIV promoter activity with BMAL1/CLOCK over-expression in HEK293T cells^19^. We and others have shown that mutation of the E-box motifs in the LTR reduced transcriptional activity^18,19^, suggesting a role for binding host factors that drive HIV-1 replication. Here, we show evidence of BMAL1 binding to all four E-box motifs in the LTR. However, this data should be interpreted with some care as the sonication protocol used in the ChIP assay can shear chromosomal DNA into ∼200-300bp fragments^42^. We previously reported that BMAL1 can bind E-box motifs in the hepatitis B viral genome^14^, supporting a model where human viruses have evolved to act in concert with host circadian transcription factors. In contrast, herpes and influenza A virus infections were enhanced in BMAL1 deficient mice^15^. These differences could result from the differential circadian modulation of viral replication in host cells vs. the anti-viral immune responses in the whole animal.

*Rorc* knock-out in mice was shown to decrease the amplitude of circadian oscillations in target genes without changing the phase or period^43^. ROR inverse agonists reduced clock gene expression in U-2 OS cells and T cells, in line with previous reports on other ROR inhibitors^30^. Importantly, we found that ROR inverse agonists reduced HIV replication and repressed the activity of LTRs from different HIV-1 clades. ROR inhibitors have been evaluated in a variety of disease and clinical models^11,44^. For instance, GSK805 was reported to reduce encephalomyelitis symptoms in a murine model of multiple sclerosis^45^ and limit intestinal inflammation^46^. GSK2981278 was evaluated in human clinical trials for the treatment of psoriasis^31,44^. These studies indicate clinical potential, but therapeutic exploitation of circadian compounds to modulate HIV-1 infection and their potential systemic effects on the host circadian rhythm requires further evaluation.

HIV-1 replication depends on a range of cellular host factors^33,34^ and we propose that many are circadian regulated. Cleret-Buhot *et al* analysed the expression of HIV-1 dependency factors between Th1 and Th17 T cells and observed increased *Rorc* and *Bmal1* transcripts in Th17 cells and an enrichment of REV-ERBα canonical pathways compared with Th1 cells^47^. A more recent report by Hiatt *et al*^35^ performed CRISPR-Cas9 silencing of previously identified HIV-host protein-protein interactions in CD4 T cells, which served as the basis for our analysis. We found that nearly half of the HIV-1 host factors are cycling in humans, however, this does not account for tissue specific gene expression. Comparing the list of HIV-1 host factors with validated BMAL1, REV-ERB and RORC target genes in murine liver or T cells identified several potential circadian regulated HIV-1 host factors. Given species and tissue specific differences in circadian gene expression^48,49^ these datasets can be further improved and reflect the limited circadian studies in human immune cells.

Bioinformatic analysis identified E-box or RORE motifs in the promoter regions of many known HIV-1 host factors, consistent with the circadian regulation of these genes. However, a more distant circadian regulation in the context of 3D-chromatin structure is possible^48^. We validated the effect of both ROR inhibitors on these genes and identified seven hits: *Csde1, Eif3m, Ranbp1, Skp1, Slirp, Spcs3* and *Timm13*. All these factors were previously identified based on their interaction with viral proteins^34,35^, and there is further evidence for their role in HIV-1 replication or circadian rhythms: The RNA binding protein CSDE1 co-localised with HIV-1 RNA in infected HeLa cells^50^ and the eukaryotic translation initiation factor 3 (EIF3) has been shown to limit HIV-1 replication^34,51^. RANBP1 is important for post transcriptional regulation of HIV-1^52^ and its silencing inhibits HIV-1 replication^53^, which is in line with our observations. As part of a ubiquitin ligase complex involved in the degradation of CRYs, SKP1 impacts cellular rhythmicity^54^. There is limited evidence for SPSC3 in regulating the cellular clock, however, it has been reported to play a role in flavivirus genesis^55^. Overall, these circadian regulated host factors provide a basis for future analysis to elucidate mechanisms defining rhythmic HIV-1 replication.

Our study mainly focused on the influence of clock factors on HIV transcription, while other stages of the viral life cycle may be influenced by circadian pathways. For instance, one of the HIV-1 entry receptors C-X-C chemokine receptor 4 (CXCR4) showed diurnal variation^56^ which could result in time-of-day dependent viral entry. We primarily investigated the influence of BMAL1, REV-ERB and ROR on HIV-1 replication, however, other circadian factors could also play a role as a Per1 short isoform was shown to inhibit HIV-1 transcription in resting CD4 T cells^57^.

Can HIV infection perturb the host circadian clock? The HIV encoded trans-activator of transcription (Tat) protein can modulate circadian rhythmicity^58,59^, however the mechanism was not defined. In contrast, Stern *et al* showed intact circadian transcriptional machinery in T cells from HIV infected subjects^20^. Clinical studies report an attenuation of diurnal blood pressure rhythms in HIV infected patients that may be due to their compromised immune status^60,61^. Pre-mature immune aging and non-AIDS related diseases increase in HIV patients with age^62^. Importantly, rhythmic activities such as sleep/wake patterns change with increasing age^63^, consistent with reports of disturbed sleep in HIV infection^64,65^. Our study shows a direct circadian regulation of HIV replication and hightlights the complex interplay between the clock, aging and sleep in HIV therapy.

Upon integration, HIV-1 establishes chronic life-long infections, and it is beneficial for the virus to adapt to the host’s physiology including its endogenous circadian rhythms. This is reflected by the balance between the host immune response and viral evasion strategies to reduce immune-associated costs. For instance, inflammatory responses in humans are reduced during the night (resting phase) when low pathogen encounter is anticipated^7^. We speculate that adaptation of the circadian regulation is advantageous for HIV as it may maximise viral replication at times when the host antiviral state is low. Our data shows that HIV-1 replication coincides with peak Bmal1 expression that aligns with a clinical study reporting high Bmal1 and HIV-1 transcript levels in CD4 T cells collected at night^20^, supporting viral adaptation to the host clock. However the impact of rhythmic anti-viral immune response remains to be defined.

Overall, our findings highlight the complexity of the virus-circadian interplay and provide fundamental insights on host pathways that regulate HIV-1 transcription. In-depth studies of circadian modifiers with antiviral properties could uncover novel drug targets which may augment existing treatment regimens for HIV and other viral infections. We hypothesise that viruses have evolved to adopt and exploit the cellular circadian machinery which strengthens the importance of future work to understand the viral-circadian interplay that will impact on our design and evaluation of new therapies.

## METHODS

### Cell lines and primary cells

HEK293T cells and human bone osteosarcoma epithelial cells (U-2 OS) were purchased from ATCC and cultured in Gibco DMEM medium (high glucose, GlutaMAX supplement, pyruvate; Life Technologies) containing 10% FBS and 1% penicillin/streptomycin (Life Technologies). U-2 OS cells stably expressing the Bmal1 promoter in front of luciferase (Bmal1-luc) were generated using the pABpuro-BlucF plasmid (Addgene #46824) and maintained in 2 µg/ml puromycin (Life Technologies). Jurkat cells were provided by Professor Xiaoning Xu (Imperial College, London, UK) and maintained in RPMI-1640 medium (Life Technologies) containing 10% FBS and 1% penicillin/streptomycin (Life Technologies).

Peripheral blood mononuclear cells (PBMCs) were isolated from leukapheresis cones purchased from NHS Blood and Transplant (Oxford, UK), with written informed consent from all donors and ethical approval for research use. CD8 T cells were depleted using a kit from Miltenyi Biotec (CD8 MicroBeads, human) and cells cultured in RPMI-1640 (Life Technologies) containing 10% FBS, 1% penicillin/streptomycin (Life Technologies), 1% sodium pyruvate (Sigma), 1% Glutamax (Life Technologies), 1% non-essential aminoacids (Life Technologies) and 2 mM beta-mercaptoethanol (Life Technologies). Cells were stimulated with 50 IU/ml IL-2 (Proleukin; Novartis), 0.01 µg/ml soluble human anti-CD3 (R & D; clone UCHT1) and 0.1 µg/ml soluble human anti-CD28 (Life Technologies; clone CD28.2) for 3 days before infection. All work with primary human cells was compliant with institutional guidelines.

### Reagents

RORC inverse agonist GSK805 (Life Technologies) and ROR inverse agonist GSK2981278 (Selleckchem) were dissolved in dimethyl sulfoxide (Life Technologies) and their cytotoxicity assessed using a lactate dehydrogenase (LDH) assay (Promega). The integrase inhibitor raltegravir was purchased from Cambridge Bioscience Ltd. Silencing RNAs for Bmal1 (ARNTL silencer select) and siRNA controls were obtained from Thermo Fisher. LIVE/Dead™ Fixable Aqua for flow-cytometry was ordered from Life Technologies.

### Plasmids

The lentiviral packaging plasmids pMD2G (#12259) and psPAX2 (#12260) were obtained from Addgene, pcDNA3.1 from Thermo Fisher and lenti-shBmal1 plasmid from ABM. The plasmid encoding NL4.3 R-E-luc was supplied by the NIBSC AIDS Repository and the VSV-G expression plasmid as previously reported^66^. HIV-LTR luciferase constructs encoding LTR regions cloned from diverse HIV-1 clades were a gift from Professor B. Paxton (University of Liverpool, UK) and were previously described^18^. The Bmal1 promoter luciferase reporter vector was purchased from Addgene (#46824). The *Bmal1* and *Clock* expression plasmids were a gift from Professor Ximing Qin (Anhui University, Hefei, China).

### Generation of viral stocks

HEK293T cells were transfected with plasmids using polyethylenimine (PEI, Polysciences). Media was replaced 4 h post transfection with DMEM media without antibiotics supplemented with 10 mM HEPES and virus harvested 48 h later. For NL4.3-luc VSV-G pseudovirus production, the NL4.3 R-E-luc and VSV-G plasmids were transfected together. To produce HIV-1 CH185 transmitted founder stocks, full-length infectious molecular clone plasmid DNA (originally obtained from Dr John Kappes, University of Alabama at Birmingham, USA^67^) was transfected using Lipofectamine (Life Technologies) or Fugene 6 (Promega) and supernatants harvested 3 days later. The viral stocks were quantified by measuring the reverse transcriptase (RT) activity using either a qPCR-based product-enhanced RT (PERT) assay^68^ or a colorimetric RT assay (Roche Life Sciences).

### Primers

Oligonucleotide sequences (Life Technologies).

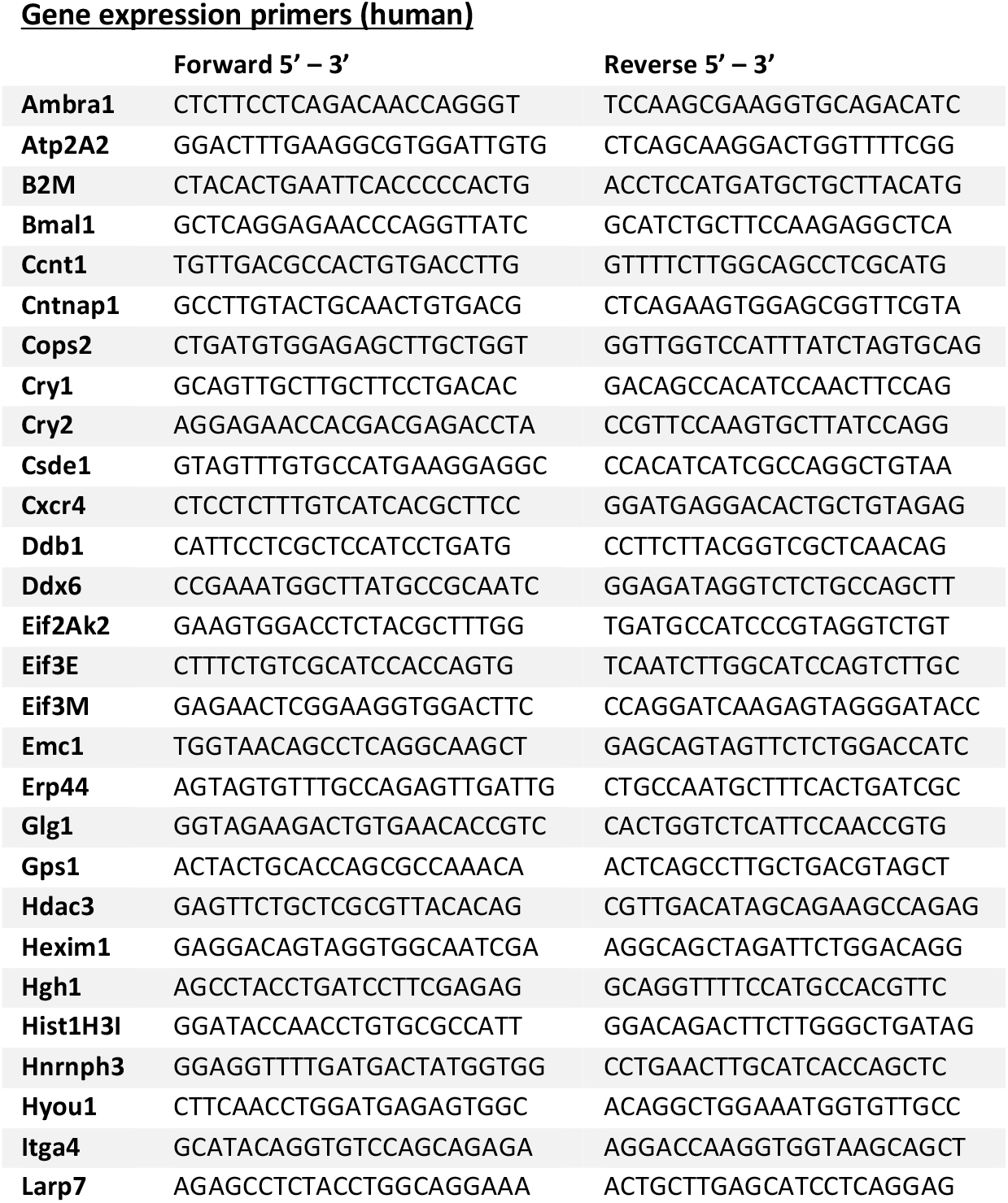
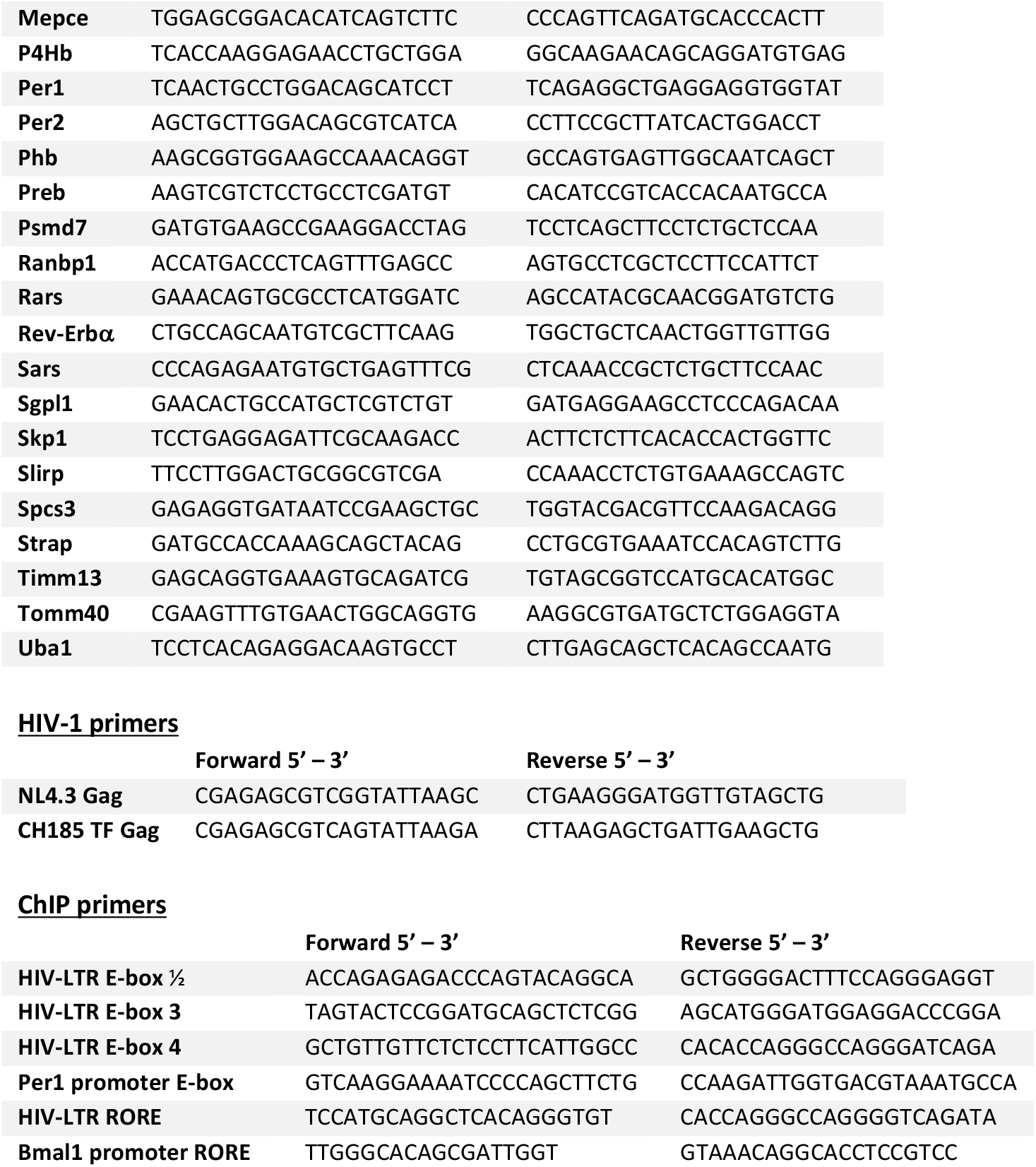

### *In vitro* HIV-1 infection experiments

To study NL4.3 R-luc VSV-G replication, U-2 OS, Jurkat cells or activated CD8 depleted PBMCs were infected with NL4.3-luc VSV-G (100 U RT/10^6^ cells) for 24 h. Cells were washed and incubated with media containing drugs at indicated concentrations for 24 h, followed by lysis for qPCR analysis or detection of luminescence using a Firefly Luciferase Assay kit (Promega) and a Glomax luminometer (Promega). To study HIV-1 replication, an infectious molecular clone (transmitted founder virus) from patient CH185 was used. Activated CD8 depleted PBMCs were spinoculated (0.25 ng RT /10^6^ cells) for 2 h and cultured in media with GSK805 for 7 days, followed by lysis and Gag RNA detection via qPCR.

### RT-qPCR

Cells were lysed and RNA extracted using an RNeasy kit (QIAGEN). For experiments with infected or plasmid transfected cells, residual DNA was digested using the TURBO DNase free kit (Thermo Fisher). Equal amounts of RNA were used for cDNA sythesis (High Capacity cDNA Kit, Applied Biosystems) and qPCR performed using Fast SYBR Master Mix (PCR Biosystems) in a LightCycler96 (Roche) with the primers listed above. mRNA expression was calculated relative to Beta-2-Microglobulin (Β2M) expression using the Ct method.

### Time course experiments

For experiments measuring luciferase expression in real time, U-2 OS cells were infected with VSV-G-pseudotyped HIV-1 NL4.3-luc (100 U RT/10^6^ cells) for 24 h or cells stably expressing a Bmal1 promoter luciferase reporter (Bmal1-luc). Cells were synchronised by serum shock with 50% FBS for 1 h, and 24 h post serum shock media was changed to DMEM lacking Phenol Red (Life Technologies), supplemented with 100 μM luciferin (VivoGlo, Promega) and each drug, respectively. Luciferase activity was measured at 30 min intervals for a period of 48 h using a CLARIOstar luminometor (BMG Labtech), with the cells kept at 37°C and 5% CO_2_. To detect rhythmic transcript levels, U-2 OS cells were infected with VSV-G-pseudotyped HIV-1 NL4.3-luc (100 U RT/10^6^ cells) for 24 h (with non-infected U-2 OS cells as control), synchronised via serum shock with 50% FBS for 1 h and from 24 h later cells harvested at 4 h intervals, followed by RNA extraction and detection by qPCR as described above. Cycling datasets were analysed with BioDare2^27^ using empirical JTK_cycle (eJTK cycle^69^) and Fast Fourier Transform Non-linear Least Squares (FFT-NLLS^26^) analysis to estimate the period, phase and amplitude of cycling transcripts. All data was normalised to control peak expression and curves fitted using Prism 9 (GraphPad). It is important to note that this normalisation sets the baseline of all data to 0. Representative raw data for each experiment can be found in **Supplementary figure 7**.

### Overexpression and silencing

*Bmal1* and *Clock* expression plasmids, or a pcDNA3.1 control, were delivered via transfection (ViaFect, Promega). For siRNA-mediated silencing, Bmal1 siRNA or a scrambled control were transfected using Fugene SI (Promega). 48 h post transfection cells were harvested for western blotting or luminescence detection using a Firefly Luciferase Assay kit (Promega) and a Glomax luminometer (Promega). To generate stable Bmal1 knock-down cells, U-2 OS cells were transduced with lentiviral vector encoding shBmal1 and cells selected using 2 μg/ml of puromycin (Life Technologies).

### Western blotting

Cells were lysed using RIPA buffer (20 mM Tris, pH 7.5, 2 mM EDTA, 150 mM NaCl, 1% NP40, and 1% sodium deoxycholate, supplemented with a protease inhibitor cocktail (Roche Complete), Laemmli sample buffer was added and samples incubated at 95°C for 5 min. Proteins were separated on a 10% polyacrylamide gel and transferred to polyvinylidene difluoride membrane (Amershan Hybond P PVDF membrane, Merck). Membranes were blocked in 5% milk in PBS/0.1% Tween-20, followed by incubation with anti-BMAL1 (Ab93806, abcam), anti-RORC (AFKJS-9, eBioscience) or anti-β-actin (A5441, Sigma) primary antibodies and appropriate HRP-conjugated secondary antibodies (DAKO). A chemiluminescence substrate (West Dura, 34076, Thermo Fisher) was used to visualise proteins using a ChemiDoc XRS+ imaging system (BioRad).

### HIV-LTR RORE and subtype LTR activity analysis

The HIV-1 5’LTR sequences deposited in the Los Alamos Database were searched in the region corresponding to the RORE (HXB2 residues 121 – 126). The program AnalyzeAlign (www.hiv.lanl.gov) was used to analyse 1266 HIV-1 sequences available in the LANL repository. To assess HIV subtype LTR activity, HIV-LTR plasmid constructs were transfected into Jurkat cells using ViaFect (Promega), 24 h post transfection cells were treated with GSK2981278 (40 μM) for 24 h and luciferase activity measured as described above.

### Chromatin immunoprecipitation (ChIP) and quantitative PCR

Jurkat cells were infected for 24 h with NL4.3-luc VSV-G (100 U RT/10^6^ cells) and then incubated in the presence of 10 μM GSK805 (or untreated control) for 24 h. Cells were fixed with 1% formaldehyde (Sigma) before quenching with 125 mM glycine. Cells were washed twice with cold PBS and lysed in SDS lysis buffer (10 mM Tris-HCl (pH 8.0), 10 mM NaCl, 1% NP-40) supplemented with a protease inhibitor cocktail (Roche Complete). Samples were diluted (1:1) with ChIP dilution buffer (0.01% SDS, 1.1% Triton, 0.2 mM EDTA; 16.7 mM Tris pH 8.1, 167 mM NaCl) and sonicated using a Bioruptor sonicator (30 min, 15 s on, 15 s off). Lysates were clarified by centrifugation and a small sample reserved as an input control in subsequent steps. Samples were precleared with Protein A agarose beads (Millipore), immunoprecipitated with anti-REV-ERBα (NRIDI) antibody (Proteintech, 14506-I-AP), anti-BMAL1 antibody (abcam, Ab3350) or rabbit IgG (Sigma, NI01) and precipitated with Protein A agarose beads. Samples were washed in low salt buffer (0.1% SDS, 1% Triton, 2 mM EDTA, 20 mM Tris pH 8.1, 150 mM NaCl), high salt buffer (0.1% SDS, 1% Triton, 2 mM EDTA, 20 mM Tris pH 8.1, 500 mM NaCl), LiCl Buffer (1% Igepal, 1 mM EDTA, 10 mM Tris pH 8.1, 250 mM LiCl, 1% sodium deoxycholate) and finally twice in TE wash buffer (10 mM Tris pH 8.0, 1 mM EDTA). Complexes were released from the beads using elution buffer (0.1 M NaHCO3, 1% SDS) and reverse crosslinked overnight at 65°C, shaking at 1400 rpm in the presence of 200 mM NaCl. After treatment with Proteinase K (Sigma) and RNaseA (Sigma), DNA was purified with MiniElute PCR Purification columns (Qiagen). qPCR was performed using a SYBR green qPCR mastermix (PCR Biosystems) in a LightCycler96 (Roche) with the primers listed above. The % input was calculated for each sample and normalised to IgG, allowing us to determine the fold enrichment of binding.

### Flow cytometry

To measure cell viability, control or drug treated cells were stained with LIVE/Dead™ Fixable Aqua (Life Technologies) and fixed with 4% PFA (Santa Cruz) for 10 min at room temperature. Sample data was acquired on a Cyan ADP flow cytometer (Beckman Coulter) and data analyzed using FlowJo (TreeStar).

### Bioinformatic analysis

The Eukaryotic Promoter Database^70^ was used to identify RORC binding motifs in circadian promoters. HIV-1 host factors were obtained from Hiatt *et al*^35^ and cycling genes identified using the Circa Database^36^ by analysing the human datasets deposited on the platform. BMAL1 regulated genes were obtained from a published ChIP-seq dataset of mouse liver^37^. REV-ERB^38^ and RORC^39^ target genes were obtained from ChIP-seq datasets of mouse T cells. Promoter regions of the genes encoding HIV-1 host factors^35^ were inspected for the presence of E-box (CANNTG, canonical CACGTG) or RORE (RGGTCA) in sequences up to −1kb upstream of the TSS using HOMER (Hypergeometric Optimization of Motif EnRichment^40^). Gene ontology (GO) analysis was performed using ShinyGO^41^, whereby each node represents an enriched GO term and related GO terms are connected by lines, whose thickness reflects the percentage of overlapping genes.

### Statistical analysis

*p* values were determined using Mann-Whitney or Kruskal-Wallis tests using Prism 9 (GraphPad). In the figures * denotes *p* < 0.05, ** *p* < 0.01, *** *p* < 0.001, all data are presented as mean values ± SEM.

## ACKNOWLEDGEMENTS

We wish to thank Prof Ximing Qin (Anhui University, Hefei, China) for *Bmal1* and *Clock* expression plasmids; Prof Bill Paxton (University of Liverpool, UK) for HIV-LTR Luc plasmids; Prof Xiaoning Xu (Imperial College, London, UK) for Jurkat cells; Dr Ye Zheng (Salk Institute, La Jolla, US) for RORC and REV-ERBα ChIP datasets; and Dr Jean-Michel Fustin (University of Manchester, UK) for his critical reading and comments on the manuscript. This work was funded by the Wellcome Trust Studentship in Infection, Immunology and Translational Medicine (HB); Wellcome Trust Award IA 200838/Z/16/Z (JAM); MRC Project Grant MR/R022011/1 (JAM); Chinese Academy of Medical Sciences Innovation Fund for Medical Science 2018-I2M-2-002 (JAM); and NIH NIAID grant UM1-AI-16456 (P Borrow).

## AUTHORS CONTRIBUTIONS

H.B. designed the study, conducted experiments and co-wrote the manuscript (MS); G.U., A.E.K., D.I. and M.S. conducted experiments; A.M., J.H.M., P. Balfe, S.V., and P. Borrow contributed to experimental design and provided comments; X.Z. designed the study and co-wrote the MS; J.A.M. designed the study and co-wrote MS.

