## Supplementary figures for "Circadian regulation of human immunodeficiency virus type 1 replication"

**Supplementary Figure 1**

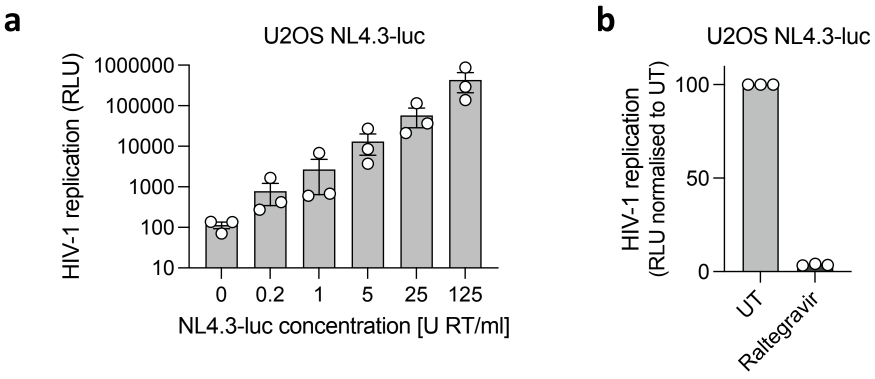


**Supplementary Figure 1. U-2 OS cells support HIV-1 infection. (a)** U-2 OS cells were infected with NL4.3-luc VSV-G for 24 h, cells lysed and luciferase activity (relative light units, RLU) measured as a read out for HIV-1 replication (mean ± S.E.M., n=3). **(b)** U-2 OS cells were pre-treated with the integrase inhibitor raltegravir (30 µM) for 24 h. Treated cells were infected with 100 U RT/ml NL4.3-luc VSV-G for 24 h in the presence of raltegravir and luciferase readings obtained 24 h post infection (mean ± S.E.M., n=3).

**Supplementary Figure 2**

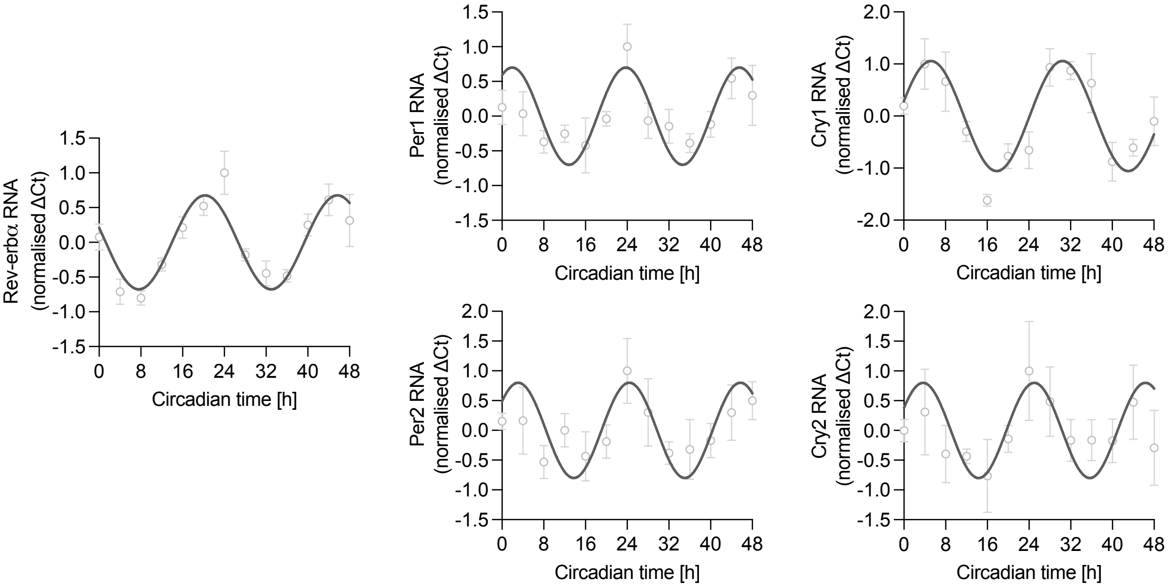


**Supplementary Figure 2. Circadian gene expression in synchronised U-2 OS cells.** U-2 OS cells were synchronised by serum shock for 1 h. 24h later, cells were harvested at 4 h intervals, RNA was extracted and expression of Rev-erbα, Per1, Per2, Cry1 and Cry2 RNAs relative to B2M housekeeper was measured by qPCR (mean ± S.E.M., n=4, normalised to peak).

### Supplementary Figure 3

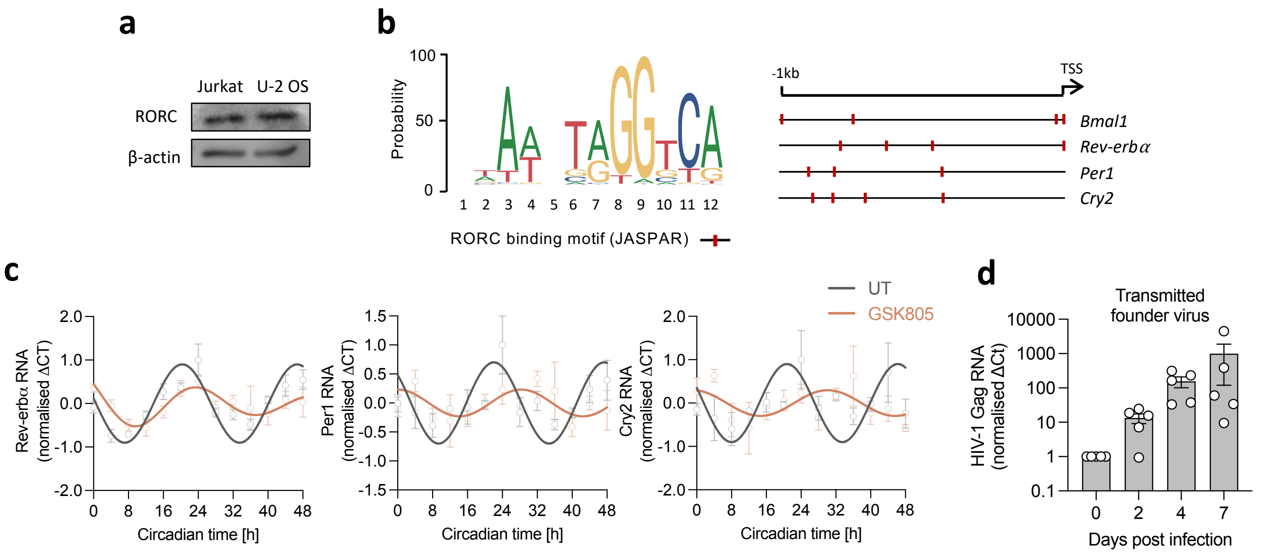


**Supplementary Figure 3. GSK805 disturbs circadian gene expression in U-2 OS and primary T cells. (a)** RORC expression in U-2 OS and Jurkat cell lysates (β-actin as control, representative of n=2). **(b)** The consensus sequence of the RORC DNA binding motif (JASPAR database) is shown, and the level of conservation is reflected by the height of the bases (y axis 0–100%). Presence and location of RORC motifs in *Bmal1, Rev-erbα, Per1* and *Cry2* promoter (1kb downstream of transcriptional start site (TSS), analysed with The Eukaryotic Promoter Database[70]) is indicated by red symbols. **(c)** U-2 OS cells were synchronised by serum shock, treated with GSK805 (10 μM) and harvested at 4 h intervals, followed by RNA extraction and qPCR detection of RNA relative to a B2M housekeeper (mean ± S.E.M., n=3, normalised to peak). **(d)** CD8 depleted PBMCs were activated for 3 days with anti-CD3/CD28 and spinoculated with transmitted founder virus CH185 for 2 h. Cells were lysed, RNA extracted, DNA digested, reverse transcribed and HIV-1 Gag RNA measured by qPCR over 7 days (mean ± S.E.M., n=5; three of the biological repeats used cells pooled from three healthy donors, two further repeats were from a single HIV-seronegative donor each).

**Supplementary Figure 4**

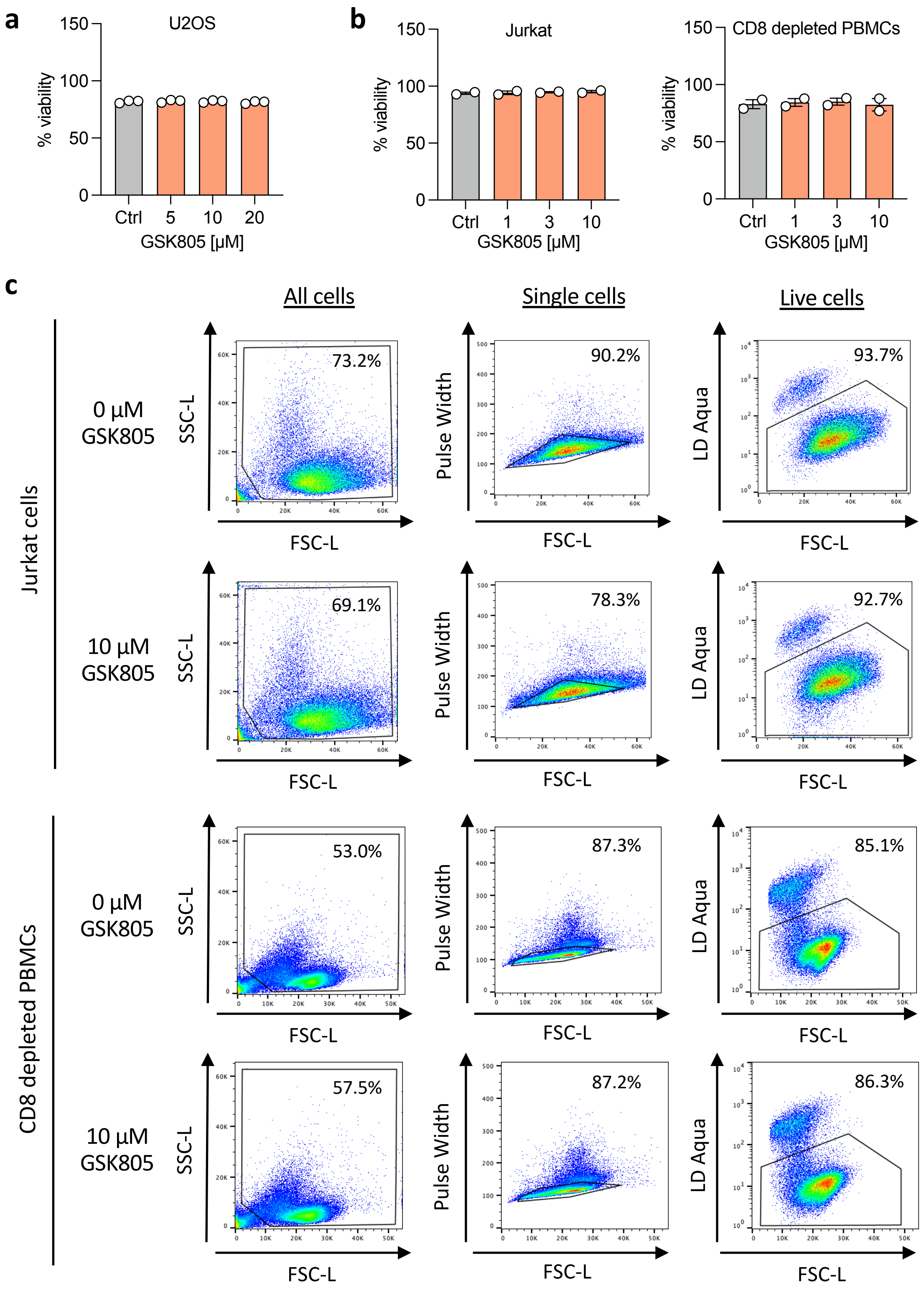


**Supplementary Figure 4. Non-cytotoxicity of GSK805 for various cell types. (a)** U-2 OS cells were treated with the RORC inverse agonist GSK805 at a range of doses for 24 h and cytotoxicity determined using an LDH assay. Data are expressed as % viability (mean ± S.E.M., n=3). **(b)** Jurkat cells or activated CD8 depleted PBMCs were treated with the ROR inverse agonist GSK805 for 24 h, and viability assessed by flow cytometry using an Aqua live-dead stain (mean + S.E.M., n=2). **(c)** Representative dot plots illustrating gating strategy and analysis. SSC = Side Scatter, FCS = Forward Scatter.

**Supplementary Figure 5**

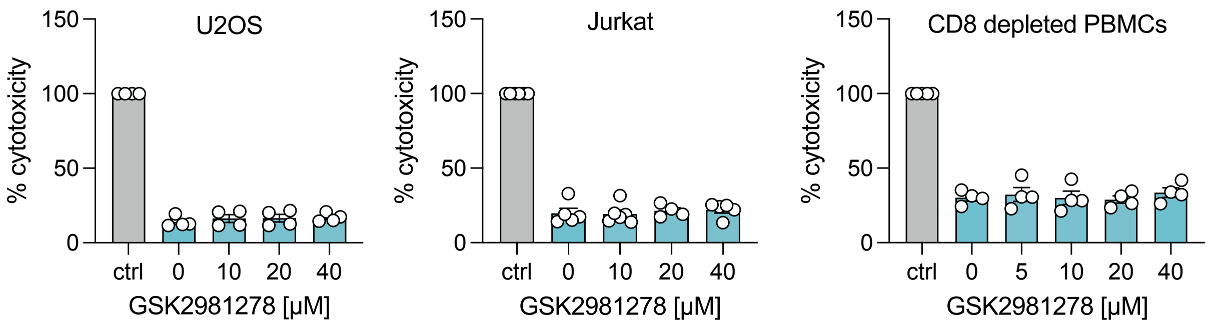


**Supplementary Figure 5. Non-cytotoxicity of GSK2981278 for various cell types.** U-2 OS, Jurkat or CD8 depleted PBMCs were treated with the ROR inverse agonist GSK2981278 at a range of doses for 24 h and cytotoxicity determined using a LDH assay (mean ± S.E.M., n=4-6). Data are expressed relative to the positive control representing total cell lysate (100% cytotoxicity).

### Supplementary Figure 6

**a**

**Cycling genes (39)**

| Dependency factors | Restriction factors |
|---|---|
| AFF1 | AMBRA1 |
| CBFB | ATP2A2 |
| CCNT1 | CEP57 |
| CDK9 | EIF2AK2 |
| CNTNAP1 | HDAC3 |
| COPS8 | HEXIM1 |
| EIF3D | LDLR |
| EIF3E | MEPCE |
| EIF3H | PELO |
| GPS1 | P4HB |
| HNRNPH3 | RANBP2 |
| MATR3 | RBM17 |
| MSH2 | SDCCAG8 |
| PARP4 | SMG6 |
| PELO | |
| PHB | |
| PREB | |
| PSIP1 | |
| RNF126 | |
| RUVBL1 | |
| TCEB1 | |
| TIMM13 | |
| TMED9 | |
| TOMM40 | |
| USP11 | |
| VDAC1 | |

**b**

**BMAL1 targets (28)**

| Dependency factors | Restriction factors |
|---|---|
| PELO | AMBRA1 |
| AFF1 | ATP2A2 |
| COPS4 | CEP57 |
| CXCR4 | DDB1 |
| EIF3E | EIF2AK2 |
| HNRNPH3 | G3BP1 |
| HYOU1 | LDLR |
| MATR3 | MEPCE |
| MCM6 | P4HB |
| NCAPD3 | PELO |
| PHB | RBM17 |
| PSMA1 | |
| RNF126 | |
| SARS | |
| SGPL1 | |
| TOMM40 | |
| VDAC1 | |
| XPO5 | |

**REV-ERB targets (31)**

| Dependency factors | Restriction factors |
|---|---|
| AFF1 | DDB1 |
| CBFB | HEXIM1 |
| CCNT1 | HUWE1 |
| CNTNAP1 | ITGA4 |
| COPS4 | LDLR |
| CSDE1 | NCOR1 |
| CXCR4 | RANBP2 |
| DDX6 | RARS |
| EIF3H | RBM17 |
| ERP44 | SDCCAG8 |
| GLG1 | SMG6 |
| ITGAL | VPRBP |
| MATR3 | |
| MCM6 | |
| MSH2 | |
| PSMA1 | |
| SPCS3 | |
| TIMM13 | |
| VDAC1 | |

**RORC targets (60)**

| Dependency factors | Restriction factors |
|---|---|
| AFF1 | AMBRA1 |
| ANKRD17 | ATP2A2 |
| CBFB | CEP57 |
| CCNT1 | COPS2 |
| CHAF1B | DDB1 |
| CHP1 | G3BP1 |
| CNTNAP1 | HEXIM1 |
| COPS3 | ITGA4 |
| COPS4 | LDLR |
| COPS8 | NCOR1 |
| CSDE1 | NDUFB10 |
| CXCR4 | RANBP2 |
| DDX6 | RBM17 |
| EIF3E | SDCCAG8 |
| EIF3H | SMG6 |
| EIF3M | VPRBP |
| ERP44 | |
| GLG1 | |
| HGH1 | |
| HYOU1 | |
| ITGAL | |
| MATR3 | |
| MCM6 | |
| MOGS | |
| MSH2 | |
| NCAPD3 | |
| NUDC | |
| PHB | |
| PHB2 | |
| PREB | |
| PSIP1 | |
| PSMA1 | |
| RUVBL1 | |
| SARS | |
| SEC61A1 | |
| SGPL1 | |
| SPCS3 | |
| STRAP | |
| TCEB1 | |
| TIMM13 | |
| TMED9 | |
| TOMM40 | |
| VDAC1 | |
| XPO5 | |

**c**

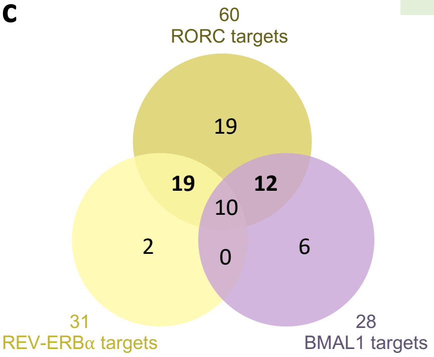


**d**

**Encoding circadian motifs in promoter (40)**

**Canonical E-box**

| Dependency factors | Restriction factors |
|---|---|
| **EMC1** | DDB1 |
| **HGH1** | EIF2AK2 |
| HNRNPH3 | **MEPCE** |
| PHB | |
| SKP1 | |
| **SPCS3** | |
| **TIMM13** | |

**RORE**

| Dependency factors | Restriction factors |
|---|---|
| CCNT1 | AMBRA1 |
| CNTNAP1 | ATP2A2 |
| CSDE1 | COPS2 |
| CXCR4 | HDAC3 |
| DDX6 | HEXIM1 |
| EIF3E | ITGA4 |
| EIF3M | LARP7 |
| **EMC1** | **MEPCE** |
| ERP44 | P4HB |
| GLG1 | RARS |
| GPS1 | |
| **HGH1** | |
| HIST1H3I | |
| HYOU1 | |
| PREB | |
| PSMD7 | |
| RANBP1 | |
| SARS | |
| SGPL1 | |
| SLIRP | |
| **SPCS3** | |
| STRAP | |
| **TIMM13** | |
| TOMM40 | |
| UBA1 | |

**Supplementary Figure 6. Cycling, BMAL1, REV-ERB and RORC regulated HIV-1 host factors. (a)** HIV-1 related host proteins (dependency and restriction factors[35]) were analysed for their rhythmic expression using the CircaDatabase[36]. **(b)** BMAL1 regulated genes[37], REV-ERB regulated genes[38] and RORC target genes[39] were compared to factors known to regulate HIV-1 replication. **(c)** Overlap of BMAL1, REV-ERB and RORC target genes. **(d)** HOMER (Hypergeometric Optimization of Motif EnRichment tool) was used to analyse promoter regions (up to -1kb from TSS) of HIV-1 host factors[35] and identified gene promoters encoding a canonical E-box motif 'CACGTG' or a ROR response element (RORE) 'RCGTCA'. Gene promoters that encode both E-box and RORE are written in bold.

**Supplementary Figure 7**

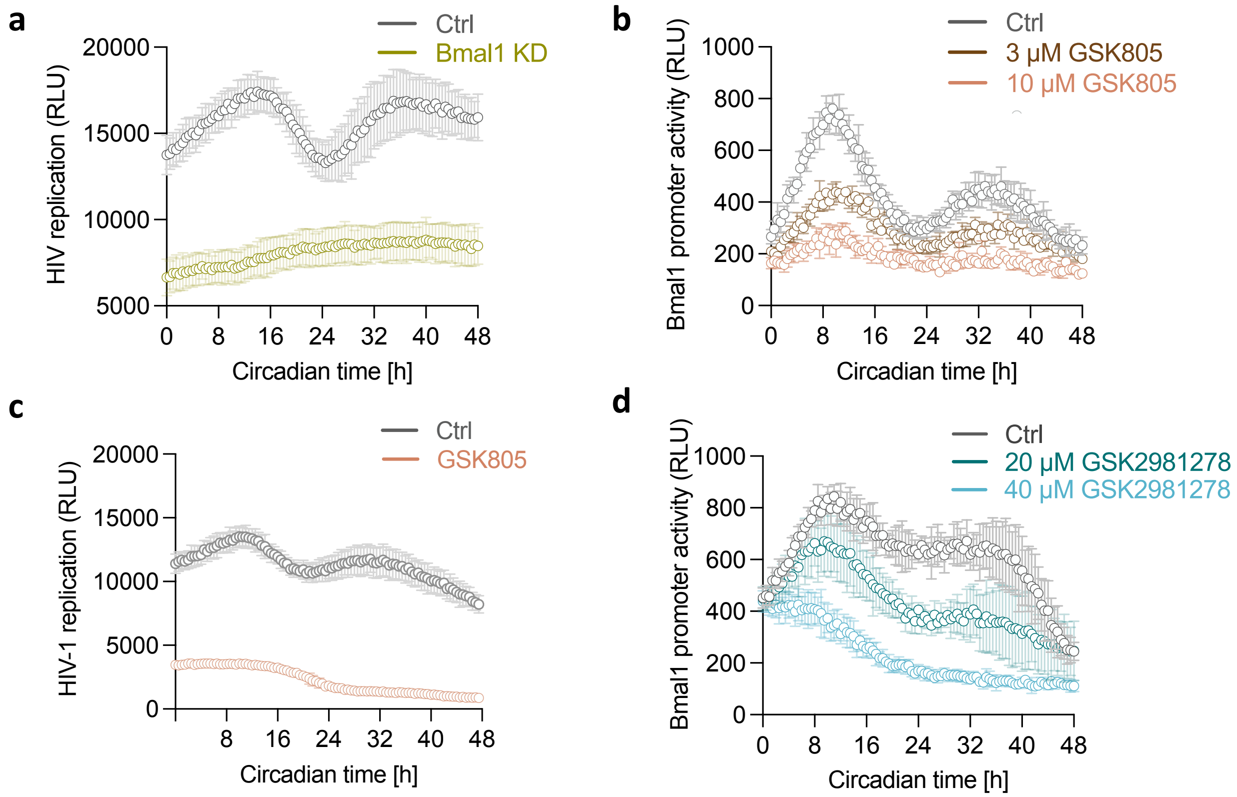


**Supplementary Figure 7. Raw luminescence values for real time measurements. (a)** U-2 OS parental control (ctrl) cells or U-2 OS Bmal1 knock-down (KD) cells generated by shRNA mediated silencing were infected with NL4.3-luc VSV-G, synchronised by serum shock and viral replication measured by luciferase readout every 30 min (representative of n=3, mean ± S.D., related to Figure 2a). **(b)** U-2 OS cells stably expressing luciferase under control of the Bmal1 promoter (Bmal1-luc) were synchronised, treated with 3 µM or 10 µM GSK805 (or untreated control) and luciferase measured at 30 min intervals (representative of n=3, mean ± S.D., related to Figure 3a). **(c)** U-2 OS cells infected with NL4.3-luc VSV-G were synchronised, treated with 10 µM GSK805 (or untreated control) and luciferase measured at 30 min intervals (representative of n=3, mean ± S.D., related to Figure 3d). **(d)** U-2 OS Bmal1-luc were synchronised, treated with 20 µM or 40 µM GSK2981278 (or untreated control) and luciferase measured at 30 min intervals (representative of n=4, mean ± S.D., related to Figure 4a).
